# SUMO2 Promotes Histone Pre-mRNA Processing by Stabilizing Histone Locus Body Interactions and Facilitating U7 snRNP Assembly

**DOI:** 10.1101/2025.02.15.638447

**Authors:** Shuying He, Pin Lyu, Marnie W. Skinner, Anthony Desotell, Brendan Foley, C.M. McCaig, Wei Wang, Jiang Qian, Liang Tong, William F. Marzluff, Michael J. Matunis

**Author notes:** Address correspondence to: Michael J. Matunis.

## Abstract

Histone mRNAs are the only non-polyadenylated mRNAs in eukaryotic cells and require specialized processing in the histone locus body (HLB), a nuclear body where essential processing factors, including the U7 snRNP, are concentrated. Recent studies have revealed that misregulation of histone pre-mRNA processing can lead to polyadenylation of histone mRNAs and disruption of histone protein homeostasis. Despite links to human disease, the factors contributing to polyadenylation of histone mRNAs and the mechanisms underlying HLB assembly and U7 snRNP biogenesis remain unclear. Here, we report novel functions of the small ubiquitin-related modifier 2 (SUMO2) in promoting histone pre-mRNA processing. Using a SUMO2 knockout osteosarcoma cell line, we identified a defect in 3’ end cleavage and a global increase in histone mRNA polyadenylation. Subsequent analysis of HLBs revealed increased dynamics and reduced levels of the U7 snRNP complex. By over-expressing U7 snRNP-specific components, Lsm11 and U7 snRNA, we rescued U7 snRNP levels and processing defects in SUMO2 knockout cells. Through analysis of Lsm11, we identified a SUMO-interacting motif in its N-terminus required for efficient formation of U7 snRNP. Collectively, we demonstrate that SUMO2 promotes histone pre-mRNA 3’ end processing by stabilizing HLB interactions and facilitating U7 snRNP assembly.

## INTRODUCTION

Small ubiquitin-related modifiers (SUMO) are posttranslational protein modifications involved in a broad range of cellular processes, including transcription, genome integrity, and cellular stress responses (Johnson 2004; Hay 2005; Vertegaal 2022). SUMO proteins can be covalently conjugated to other proteins through an E1-E2-E3 enzyme cascade, preferentially onto lysines within a ψKX(D/E) consensus motif (ψ, hydrophobic amino acids). Conjugation is reversible by SUMO-specific isopeptidases (SENPs) (Johnson 2004; Claessens and Vertegaal 2024). In addition, SUMO interacts non-covalently with proteins containing SUMO-interacting motifs (SIMs), characterized by a hydrophobic core of four amino acids adjacent to negatively charged residues (Hecker et al. 2006; Yau et al. 2021). Mammalian cells express multiple SUMO paralogs, with SUMO1, SUMO2, and SUMO3 being the best characterized. SUMO2 and SUMO3 share 97% sequence identity and are referred to as SUMO2/3. In contrast, SUMO1 and SUMO2/3 share 45% sequence identity (Saitoh and Hinchey 2000; Wang and Matunis 2023). SUMO2/3 also contains an N-terminal consensus motif (centered around K11) that allows for more efficient formation of polymeric chains compared to SUMO1 (Tatham et al. 2001). These findings suggest that SUMO paralogs have distinct functions. Consistent with this, RNA-Seq analysis revealed significantly different patterns of paralog expression across normal human tissues (Bouchard et al. 2021). To explore SUMO paralog-specific functions, we previously generated SUMO1 and SUMO2 knockout osteosarcoma (U2OS) cell lines. SUMO3 is expressed at exceedingly low levels in U2OS cells, allowing us to systematically identify and characterize paralog-specific functions of both SUMO1 and SUMO2 in various cellular processes (Bouchard et al. 2021; Wang et al. 2023). Here, using these cell lines, we report a unique function of SUMO2 in promoting histone pre-mRNA processing.

Replication-dependent histone genes, encoding the only non-polyadenylated mRNAs in eukaryotic cells, are arranged in genomic clusters, with a large cluster (*HIST1*) on chromosome 6 and a smaller cluster (*HIST2*) on chromosome 1 (Marzluff et al. 2002; Duronio and Marzluff 2017). These gene clusters are assembled into membraneless nuclear organelles known as histone locus bodies (HLBs) that contain transcription and processing factors required for histone mRNA expression. Specifically, the nuclear protein ataxia-telangiectasia locus (NPAT) and FLICE-associated huge protein (FLASH) are concentrated in HLBs and are required for HLB assembly and histone gene activation and pre-mRNA processing, respectively (Ma et al. 2000; Barcaroli et al. 2006; Yang et al. 2009). Histone genes lack introns, and thus the only processing required to form histone mRNAs is a 3’ endonucleolytic cleavage reaction (Fig 1A) (Pandey et al. 1990). Stem-loop binding protein (SLBP) binds to the stem loop on histone pre-mRNAs and stabilizes the base pairing of U7 snRNA to the 3’ histone downstream element (Fig 1B) (Dominski et al. 1999). FLASH interacts with Lsm11 in the U7 snRNP, forming an interface that recruits the histone cleavage complex that cleaves histone pre-mRNAs (Yang et al. 2009; Yang et al. 2013).

**Figure 1.**
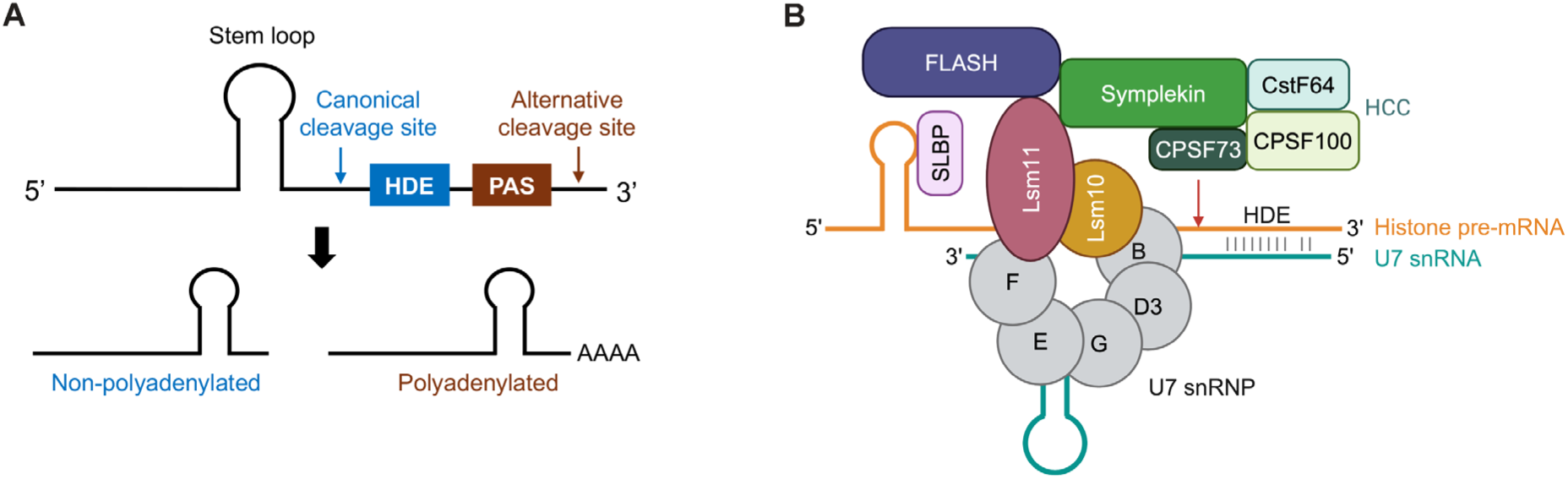
Histone pre-mRNA processing pathways and factors. (A) A schematic showing the canonical and alternative histone pre-mRNA processing pathways. HDE = histone downstream element. PAS = polyadenylation signal. (B) A schematic of the U7 snRNP complex and histone cleavage complex (HCC) involved in canonical histone pre-mRNA processing (created with BioRender.com).

The U7 snRNP and associated factors constitute a multisubunit RNA-guided endonuclease (Fig 1B) (Sun et al. 2020). The U7 snRNP itself consists of a U7 snRNA and a ring structure of seven Sm proteins. In the U7 Sm ring, five Sm proteins, B, D3, E, F, G, are shared with spliceosomal snRNPs, while SmD1 and SmD2 are replaced by U7-specific proteins, Lsm10 and Lsm11 (Pillai et al. 2001; Pillai et al. 2003). Lsm11 contains a long N-terminal tail that mediates the interaction of U7 snRNP with FLASH (Yang et al. 2009; Burch et al. 2011). In cells, histone pre-mRNA processing is tightly coupled with transcription termination and reduction in the concentration of processing factors, including U7 snRNP, disrupts this coupling (Lanzotti et al. 2002; Yang et al. 2009; Burch et al. 2011; Tatomer et al. 2016). Uncoupling results in readthrough past the normal termination site and alternative processing and polyadenylation of the subset of histone gene mRNAs that contain downstream polyadenylation sites (Fig 1A) (Wagner et al. 2007; Lyons et al. 2016).

Though misregulation of histone pre-mRNA processing typically results in polyadenylation of only 2-3% of total expressed histone mRNAs, these changes affect histone protein expression levels and cell function (Narita et al. 2007; Kohn et al. 2015). Increased polyadenylation results in the continued presence of histone mRNAs and expression of histone proteins outside of S phase, thus disrupting replication-dependent regulation (Harris et al. 1991; Whitfield et al. 2000; Brocato et al. 2014). Patients with Aicardi-Goutières syndrome carry mutations in the *LSM11* and *RNU7-1* (encoding U7 snRNA) genes. These mutations result in increased histone mRNA polyadenylation, which disrupts histone protein homeostasis and chromatin structure and results in activation of type I interferon signaling (Uggenti et al. 2020). Other studies have shown that in spinal muscular atrophy models, survival motor neuron (SMN) protein dysfunction results in U7 snRNP assembly defects, increased histone mRNA polyadenylation, and altered histone protein levels. These defects may contribute to neuromuscular pathology (Tisdale et al. 2022).

Despite the implication of U7 snRNP in disease, mechanisms governing U7 snRNP assembly and targeting to HLBs remain elusive. The SMN complex, composed of SMN, Gemin2-8 and Unrip, mediates the assembly of Sm rings onto both spliceosomal snRNAs and U7 snRNA (Battle et al. 2006a). However, U7 snRNP assembly is unique in not requiring Gemin5, which specifically recognizes spliceosomal snRNAs (Battle et al. 2006b). Specificity factors required for U7 snRNP assembly remain unknown.

Here, we report reduced canonical histone pre-mRNA 3’ end cleavage and increased polyadenylation of histone mRNAs in SUMO2 knockout cells. We demonstrate novel functions of SUMO2 in promoting histone pre-mRNA processing through two facets. First, SUMO2 stabilizes interactions of FLASH within HLBs. Second, SUMO2 promotes U7 snRNP assembly and expression levels through interactions with a SIM in the N-terminus of Lsm11. Taken together, our findings reveal SUMO2 as an important component of the canonical histone pre-mRNA processing pathway.

## RESULTS

### Paralog-specific functions of SUMO2 in promoting histone pre-mRNA processing

We previously generated U2OS SUMO1 knockout (S1KO) and SUMO2 knockout (S2KO) cell lines with CRISPR-Cas9, as well as SUMO2 knockout rescue cell lines (S2R) by stable SUMO2 re-expression (Bouchard et al. 2021). Through RNA-Seq analysis of poly(A)-selected mRNAs, we identified 20 replication-dependent histone genes with apparent increases in gene expression in S2KO cells compared to WT cells (Fig 2A). In contrast, the expression of these histone genes remained unchanged or downregulated in S1KO cells. Similar changes were not observed with replication-independent histone genes (Fig S1A). RT-qPCR analysis using oligo-dT primers of selected replication-dependent histone mRNAs, including *H1-2*, *H2BC21*, and *H4C8* as previously shown (Bouchard et al. 2021), and *H2AC6* as shown here (Fig S1B), validated this result and revealed that this increase was reversed by stable re-expression of SUMO2.

**Figure 2.**
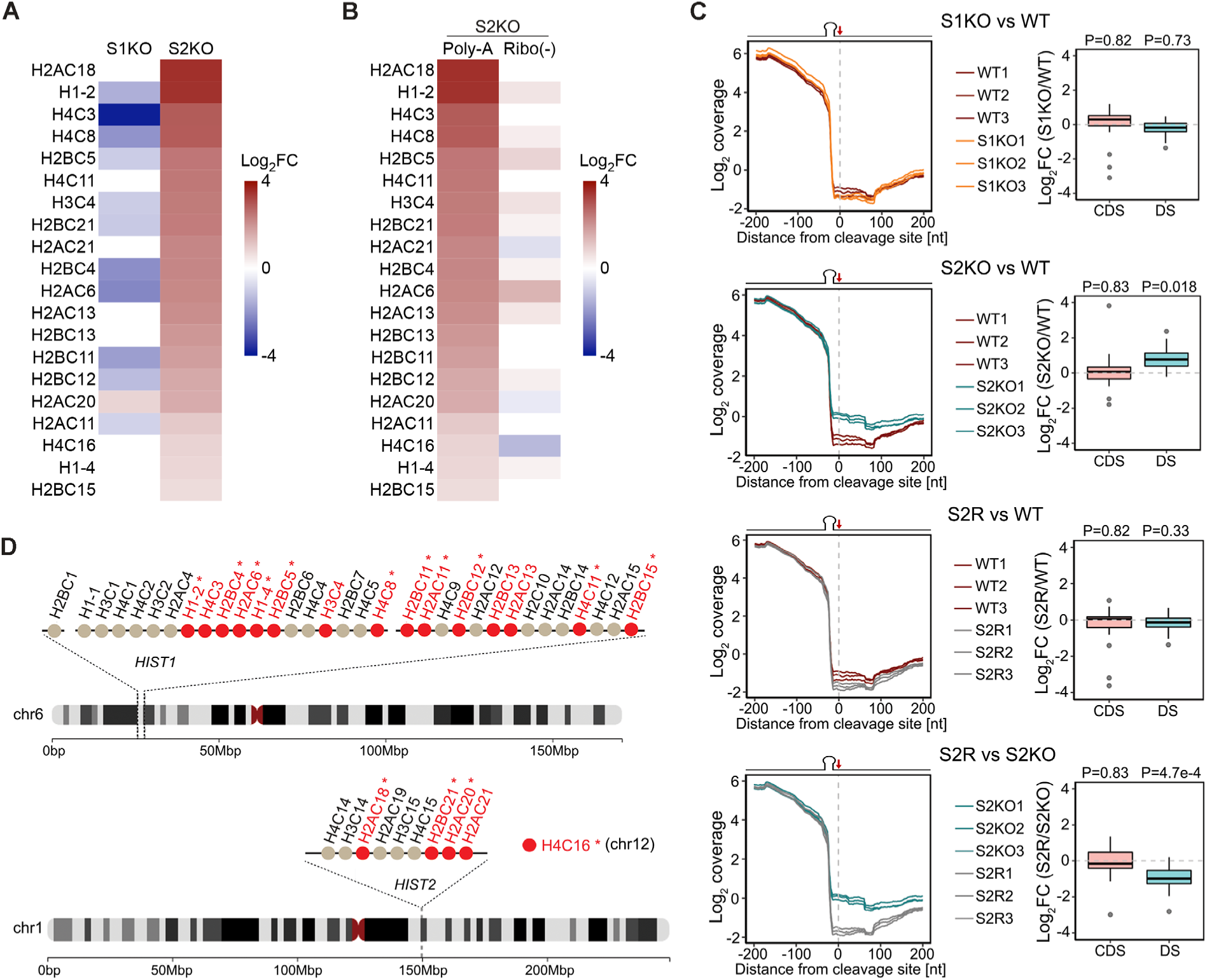
SUMO2 functions in histone pre-mRNA 3’ end processing. (A) A heatmap showing histone genes with increased mRNA polyadenylation revealed by poly(A) RNA-Seq in S1KO and S2KO cells, compared to U2OS WT cells (Log_2_FC = log_2_ fold change). (B) A heatmap comparing histone mRNA polyadenylation (measured by poly(A) RNA-Seq) to total histone gene expression (measured by ribo-minus RNA-Seq) in S2KO cells compared to WT cells. (C) Line graphs (left panels) showing combined ribo-minus RNA-Seq signals upstream and downstream of the canonical cleavage site in 43 expressed replication-dependent histone genes. The steep decrease in coverage ∼20 bp upstream of the cleavage site is due to inefficient priming of the mRNA stem loop (Welch et al. 2015). Box plots (right panels) comparing changes in ribo-minus RNA-Seq reads in coding sequences (CDS) and sequences downstream of the canonical cleavage site (DS) in U2OS cells. Three biological replicates were analyzed for each cell line. (D) A schematic of the 43 replication-dependent histone genes analyzed in panel C and their locations within histone clusters *HIST1* and *HIST2* on chromosomes 6 and 1. *H4C16* is on chromosome 12. Histone genes with increased mRNA polyadenylation in S2KO cells are highlighted in red. Histone mRNAs previously reported to be polyadenylated under disease conditions are indicated by an “*”. *P* values were calculated using two-way ANOVA tests.

Given that replication-dependent histone mRNAs are normally not polyadenylated, we hypothesized that this apparent increase in gene expression could result from defects in histone pre-mRNA processing and increased polyadenylation. To test this hypothesis, we performed RNA-Seq analysis using ribo-minus selected mRNAs that capture total (non-polyadenylated + polyadenylated) histone mRNA levels. Statistical analysis revealed consistency across three biological RNA-Seq replicates (Fig S1C). In addition, differential gene expression analysis showed a negative correlation between S2KO and S2R cells, indicating a global rescue of gene expression (Fig S1D). In comparison to poly(A) RNA-Seq analysis, ribo-minus RNA-Seq revealed only minor changes in total histone mRNA expression levels in S2KO cells compared to WT cells, indicating that the apparent increase was due to increased polyadenylation (Fig 2B).

To further investigate alterations in histone pre-mRNA 3’ end processing, we analyzed ribo-minus RNA-Seq reads at the 3’ ends of 43 replication-dependent histone gene mRNAs expressed in U2OS cells (Fig 2C). We observed a collective 2-fold increase in levels of mis-processed histone mRNA transcripts in S2KO cells compared to WT cells. This defect in 3’ end processing was rescued by stable re-expression of SUMO2, and no obvious changes were detected in S1KO cells. To assess if mis-processing was specific to replication-dependent histone genes, we analyzed 3’ end cleavage of 2,000 randomly selected genes expressed in U2OS cells and observed no changes between S2KO and WT cells (Fig S1E).

Defects in 3’ end cleavage of replication-dependent histone mRNAs in S2KO cells is further illustrated by representative analysis of histone *H1-2* ribo-minus RNA-Seq reads (Fig S1F). An average of ∼10,000 reads were mapped to the coding sequences of both WT and S2KO cells, suggesting no major changes in total gene expression. In contrast, the numbers of reads mapped downstream of the cleavage site increased from ∼100 in WT cells to ∼400 in S2KO cells, suggesting a 4-fold increase in mis-processed transcripts. These RNA-Seq reads showed that mis-processed *H1-2* mRNAs represented ∼1% of total expressed *H1-2* mRNAs in WT cells and ∼4% in S2KO cells. This is consistent with previously reported histone pre-mRNA processing defects where mis-processed mRNAs constitute a small fraction of total expressed mRNAs (Narita et al. 2007; Kohn et al. 2015). An increase in mis-processed mRNAs was not observed in S1KO cells and was rescued by stable re-expression of SUMO2.

The histone genes whose mRNAs exhibited a defect in cleavage and increased polyadenylation are distributed throughout the *HIST1* and *HIST2* histone gene clusters (Fig 2D). mRNAs affected also included those for *H4C16*, a replication-dependent histone gene located on chromosome 12 (Marzluff et al. 2002). Notably, a significant fraction of these genes (15/20) corresponds to histone genes whose 3’ end cleavage and polyadenylation is similarly altered in disease conditions and differentiated cells (Kari et al. 2013; Lyons et al. 2016; Uggenti et al. 2020). Together, these results reveal paralog-specific functions of SUMO2 in promoting the efficiency of canonical 3’ end processing of replication-dependent histone pre-mRNAs.

### SUMO2 stabilizes FLASH dynamics

Efficient histone mRNA biogenesis requires concentration of 3’ end processing factors in HLBs (Tatomer et al. 2016). Previous studies have identified sumoylation as a key regulator of promyelocytic leukemia nuclear body (PML-NB) assembly and function, raising the question of whether sumoylation may have similar roles in HLBs (Banani et al. 2016; Lallemand-Breitenbach and de The 2018). Multiple HLB-associated factors have been identified as SUMO substrates, including FLASH, which also harbors tandem SIMs (Vethantham et al. 2007; Alm-Kristiansen et al. 2009; Sun and Hunter 2012). We analyzed the localization of FLASH and another core HLB-associated protein, NPAT, by immunofluorescence microscopy and observed no defects in HLB formation in S2KO cells (Fig 3A). We then analyzed HLB dynamic interactions through live-cell microscopy of GFP-tagged miniFLASH. miniFLASH is a fusion of amino acids 1-200 and 1751-1982 of FLASH that has previously been used as a reporter for HLB dynamics (Kohn et al. 2015). miniFLASH retains a previously reported SUMO modification site and SIM (Alm-Kristiansen et al. 2009; Sun and Hunter 2012). We analyzed the association of GFP-miniFLASH with HLBs using fluorescence recovery after photobleaching (FRAP) analysis and observed an ∼2-fold increase in the mobile fraction in S2KO cells compared to WT cells based on the percent recovery, suggesting an increase in FLASH dynamic interactions within HLBs in the absence of SUMO2 (Fig 3B). We also analyzed the number of HLB foci and HLB morphology by immunofluorescence microscopy with NPAT antibodies. Consistent with a role for SUMO2 in HLBs, we observed a decrease in the number of HLB foci and an increase in their size in S2KO cells compared to WT cells. No change in NPAT signal intensity was detected (Fig 3C). These changes in FLASH dynamics and HLB morphology were partially rescued by stable SUMO2 re-expression.

**Figure 3.**
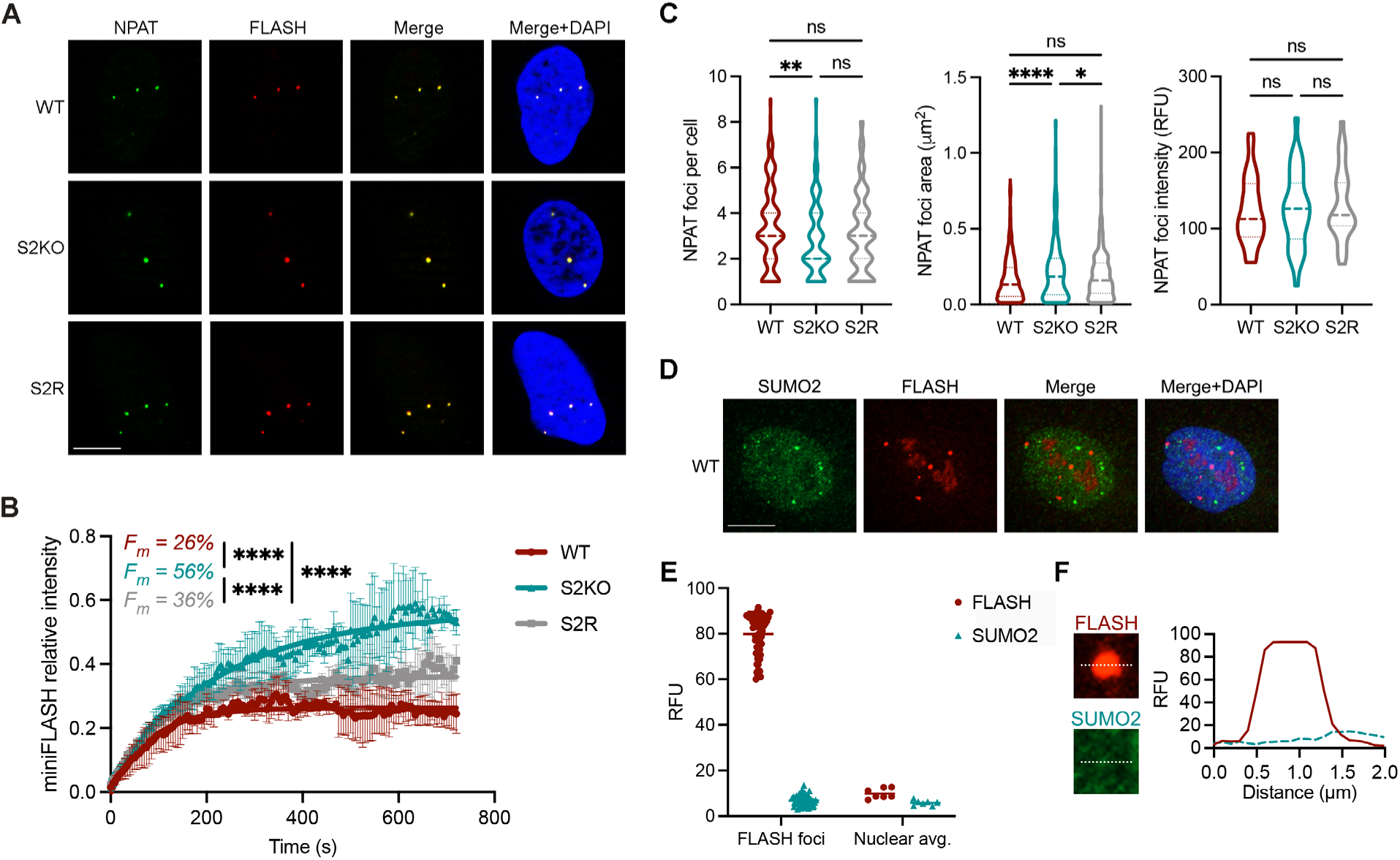
SUMO2 effects on HLBs. (A) Immunofluorescence microscopy of U2OS cells with antibodies against NPAT and FLASH. (B) FRAP analysis of GFP-miniFLASH expressed in WT, S2KO and S2R cells. An average of 30 HLBs were analyzed for each cell line from three independent replicates. F_m_ = mobile fraction. (C) Quantification of NPAT foci number per cell, area (μm^2^), and fluorescence intensity. An average of 180 cells were analyzed for each cell line from three independent replicates. (D) Immunofluorescence microscopy with antibodies against SUMO2 and FLASH in WT cells. The SUMO2 foci are likely PML-NBs. (E) Quantification of FLASH and SUMO2 fluorescence intensity within FLASH foci and the nuclear average. Signals from 7 cells were analyzed. (F) Co-localization plot of FLASH and SUMO2 signals. *P* values were calculated using one-way ANOVA tests (ns = not significant, * *p* < 0.05, ** *p* < 0.01, *** *p* < 0.001, **** *p* < 0.0001). Scale bar = 10 µm. RFU = relative fluorescence unit.

To further understand the role of SUMO2 in HLBs, we investigated whether it is enriched in them, similar to its enrichment in PML-NBs (Lallemand-Breitenbach and de The 2018). Using confocal immunofluorescence microscopy and antibodies against FLASH and SUMO2, we observed no obvious enrichment of SUMO2 within HLBs (Fig 3D). Quantification revealed that SUMO2 signal within HLBs was comparable to its nuclear average, indicating that SUMO2 is neither concentrated, nor excluded, from HLBs (Fig 3E, F). We also tested whether HLB-associated proteins are modified by SUMO2. GFP-miniFLASH was transiently expressed in WT cells and immunopurified using GFP-trap beads followed by western blotting with antibodies against SUMO2 (Fig S2). High molecular weight SUMO2 signal was detected in the pull-down fraction of GFP-miniFLASH, suggesting that FLASH and/or FLASH-associated proteins are modified by SUMO2. Together, these results demonstrate that SUMO2 stabilizes FLASH interactions in HLBs, potentially through modification of FLASH and/or FLASH-associated proteins.

### SUMO2 modulates U7 snRNP levels

Altered FLASH dynamics suggests potential changes in the localization or concentration of key histone pre-mRNA processing factors in HLBs. Using immunofluorescence microscopy, we analyzed the localization of the U7 snRNP complex with antibodies against Lsm10 and Lsm11 and found that detection of both proteins was significantly reduced in HLBs of S2KO cells (Fig 4A, B). Detection of Lsm10 in HLBs was reduced from 80% of WT cells to 30% of S2KO cells (Fig 4C). Similarly, detection of Lsm11 in HLBs was reduced from 60% of WT cells to 10% of S2KO cells (Fig 4D). This defect in U7 snRNP localization was rescued by stable SUMO2 re-expression.

**Figure 4.**
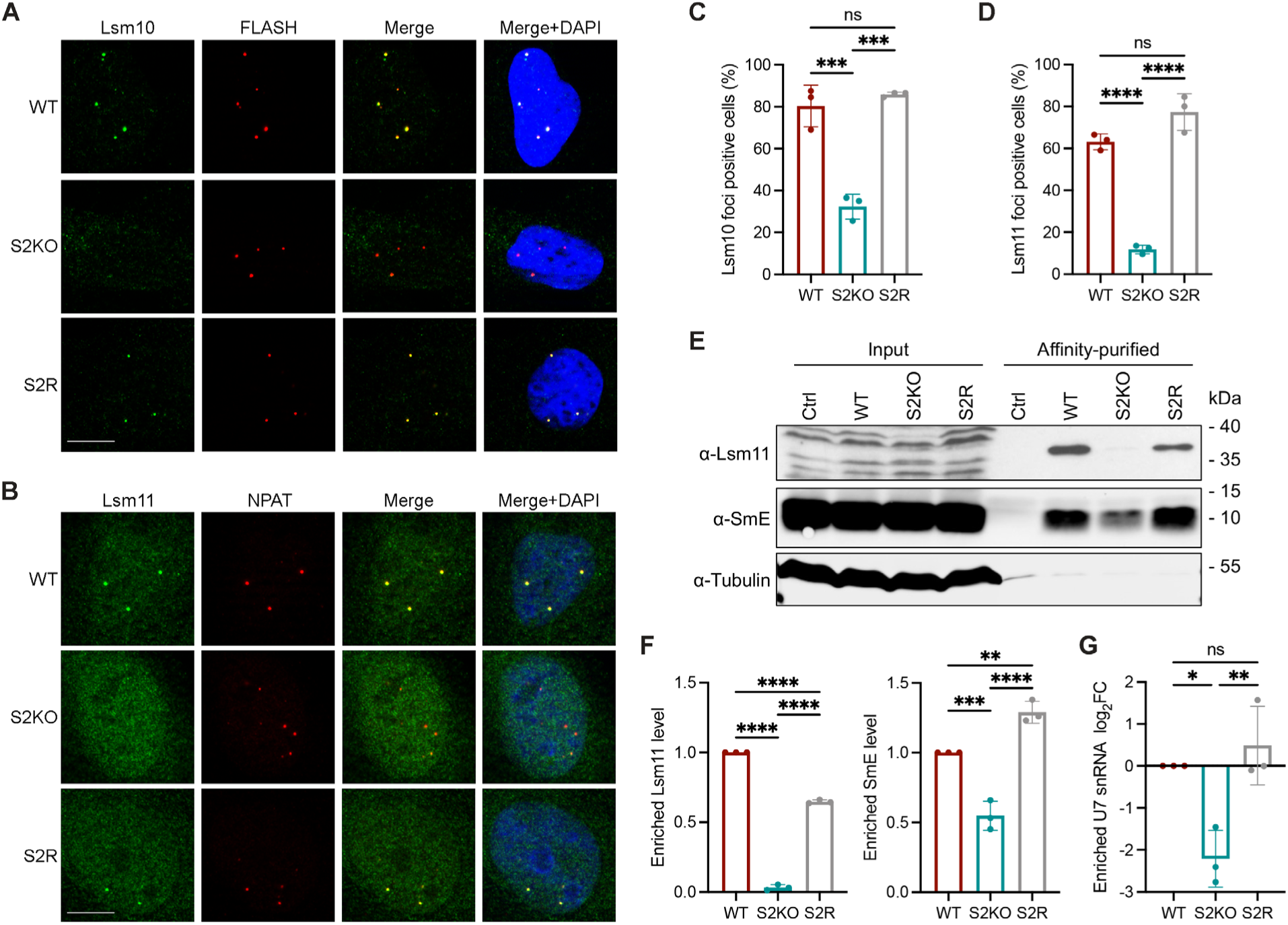
SUMO2 modulates U7 snRNP levels. Immunofluorescence microscopy with (A) antibodies against Lsm10 and FLASH or (B) antibodies against Lsm11 and NPAT. Scale bar = 10 µm. Quantification of the percentage of cells with (C) Lsm10 or (D) Lsm11 detectable in HLBs. An average of 600 cells were analyzed for each cell line from three independent replicates. (E) Western blot analysis of affinity-purified U7 snRNP complexes using U7 snRNA antisense oligos. Control (Ctrl) condition was WT cell lysate incubated with scramble oligos. Bands detected by Lsm11 antibodies in input fractions corresponded to cross-reacting proteins. (F) Quantification of enriched Lsm11 and SmE levels from three independent replicates. (G) Quantification of enriched U7 snRNA levels measured by RT-qPCR from three independent replicates. *P* values were calculated using one-way ANOVA tests (ns = not significant, * *p* < 0.05, ** *p* < 0.01, *** *p* < 0.001, **** *p* < 0.0001).

Defects in U7 snRNP concentration in HLBs could result from either reduced U7 snRNP expression or impaired association with HLBs. We first examined U7 snRNP expression levels using affinity purification followed by western blotting and RT-qPCR analysis. U7 snRNP complexes were purified from WT and S2KO whole cell lysates using biotin-tagged antisense oligos complementary to the U7 snRNA or scramble oligos as control. Antibodies against Lsm11 and SmE were used to measure levels of purified complexes. A significant decrease in U7 snRNP levels was observed in S2KO cells, as evidenced by an ∼50% reduction in SmE signal and 95% reduction in Lsm11 signal in the pull-down fraction (Fig 4E, F). Differences in levels of SmE compared to Lsm11 likely result from co-purification of other highly abundant spliceosomal snRNPs which also contain the SmE protein (Smith et al. 1991). We next extracted U7 snRNA from purified complexes and measured its levels by RT-qPCR. This analysis revealed an 80% decrease of U7 snRNA in S2KO cells compared to WT cells, consistent with the reduction in Lsm11 levels (Fig 4G). This reduction in U7 snRNP expression levels was partially rescued by SUMO2 re-expression based on Lsm11 levels. These results indicate that defects in U7 snRNP concentration within HLBs are due to reduced U7 snRNP levels.

### Rescue of U7 snRNP levels requires over-expression of Lsm11 and U7 snRNA

U7 snRNP is maintained at low levels in cells compared to the spliceosomal snRNPs (U1-U6), due in part to the low expression of Lsm10 and Lsm11 (Grimm et al. 1993; Pillai et al. 2001; Pillai et al. 2003). Efforts to directly quantify Lsm10 and Lsm11 protein levels by western blotting of whole cell lysates were unsuccessful due to their low expression levels. We therefore turned to rescue experiments involving over-expression of Lsm10, Lsm11, and U7 snRNA.

We constructed mammalian expression plasmids for expressing V5-tagged Lsm10 and Lsm11 individually, as well as a plasmid for co-expression, as previously reported (Fig S3A) (Tisdale et al. 2022). U7 snRNA was generated using in vitro transcription and capping as previously described (Boskovic et al. 2020). Cells were transiently transfected with these plasmids individually, or in combination with U7 snRNA, and U7 snRNP expression levels were monitored as described above. Transfection of U7 snRNA and Lsm10, either individually or combined, resulted in no significant rescue of U7 snRNP levels in S2KO cells compared to WT cells under the same conditions (Fig 5A, B). Transfection of Lsm11 alone resulted in an ∼25% rescue of U7 snRNP levels, similar to co-expression of Lsm11 and Lsm10 (Figure 5C, D, S3A). Co-transfection of Lsm11 and U7 snRNA resulted in an ∼60% rescue of U7 snRNP levels, comparable to co-transfection of Lsm10, Lsm11, and U7 snRNA (Fig 5C, D). Thus, maximal rescue of U7 snRNP expression levels required increased expression of both Lsm11 and U7 snRNA (Fig 5E).

**Figure 5.**
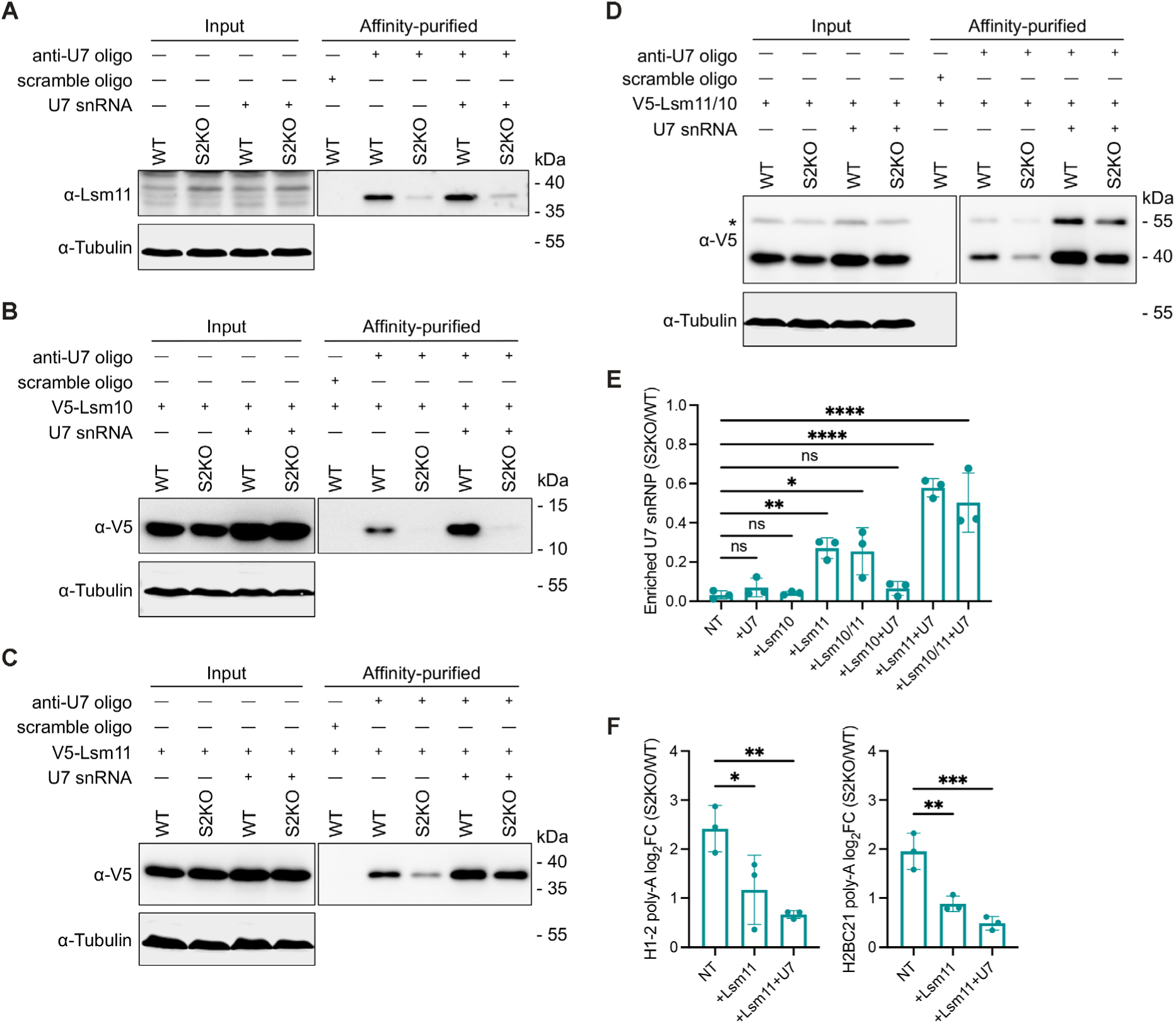
Rescue of U7 snRNP levels requires over-expression of Lsm11 and U7 snRNA. Western blot analysis of U7 snRNP complexes affinity-purified from cells transfected with (A) U7 snRNA, (B) V5-Lsm10 individually or in combination with U7 snRNA, (C) V5-Lsm11 individually or in combination with U7 snRNA, (D) V5-Lsm11/Lsm10 individually or in combination with U7 snRNA. “*” indicates uncleaved Lsm10/11 fusion protein. (E) Quantification of enriched U7 snRNP complex levels from S2KO cells relative to WT cells over-expressing the indicated factors. NT = not transfected. U7 = U7 snRNA. Three independent replicates were analyzed for each condition. (F) RT-qPCR analysis of relative polyadenylated *H1-2* and *H2BC21* mRNA levels in S2KO cells, S2KO cells stably expressing V5-Lsm11 (+Lsm11), or S2KO cells stably expressing V5-Lsm11 and transfected with U7 snRNA (+Lsm11+U7), compared to WT cells (log_2_FC = 0) under the same conditions. Three independent replicates were analyzed for each condition. *P* values were calculated using one-way ANOVA tests (ns = not significant, * *p* < 0.05, ** *p* < 0.01, *** *p* < 0.001, **** *p* < 0.0001).

We next investigated whether rescue of U7 snRNP expression levels by increased Lsm11 and U7 snRNA expression also rescued histone pre-mRNA processing and polyadenylation defects in S2KO cells. For these experiments, we generated U2OS WT and S2KO cell lines stably expressing V5-tagged Lsm11. As observed in the above transient transfection assays, U7 snRNP levels were partially rescued in these Lsm11 over-expressing cells, and the level of rescue was enhanced by U7 snRNA transfection (Fig S3B). Consistent with this effect of Lsm11 over-expression, the percentage of cells containing visible Lsm11 labeled HLBs increased from 10% in S2KO cells to 30% in S2KO cells stably expressing V5-Lsm11, and this percentage was unchanged in WT cells stably expressing V5-Lsm11 compared to WT cells (Fig S3C, 4D). Using these stable cell lines, we transiently transfected U7 snRNA and measured polyadenylation levels of two of the histone mRNAs affected in S2KO cells (*H1-2* and *H2BC21*) using RT-qPCR (Fig 5F). Polyadenylation of both *H1-2* and *H2BC21* mRNAs was ∼4-5 fold higher in S2KO cells compared to WT cells. This difference was reduced to ∼2 fold in S2KO cells stably expressing Lsm11 and further reduced to ∼1.4 fold by co-expressing U7 snRNA, compared to WT cells under the same condition. These results reveal that reduced U7 snRNP levels is a major cause of histone pre-mRNA processing defects in S2KO cells.

### SUMO2 affects U7 snRNP levels independent of core SMN complex function

Sumoylation is critical for the assembly, function, and localization of the SMN complex (Riboldi et al. 2021). To test whether reduced U7 snRNP levels are due to defects in SMN complex assembly or function, we measured spliceosomal snRNA levels in whole cell extracts using RT-qPCR (Fig 6A). While U1, U2, U4, U6, U11, U12, U4atac, and U6atac snRNAs remained unaffected, U5 snRNA levels were significantly reduced in S2KO cells. To further validate the reductions in U5 snRNP and U7 snRNP levels, we immunopurified spliceosomal snRNPs and U7 snRNP from whole cell lysates with the Y12 antibody that recognizes Sm proteins B and D, followed by RNA extraction and RT-qPCR analysis (Fig 6B) (Hirakata et al. 1993). Using U1 snRNP levels as control, we confirmed that both U5 snRNP and U7 snRNP levels were downregulated in S2KO cells and partially rescued by stable SUMO2 re-expression. All other snRNPs were unchanged.

**Figure 6.**
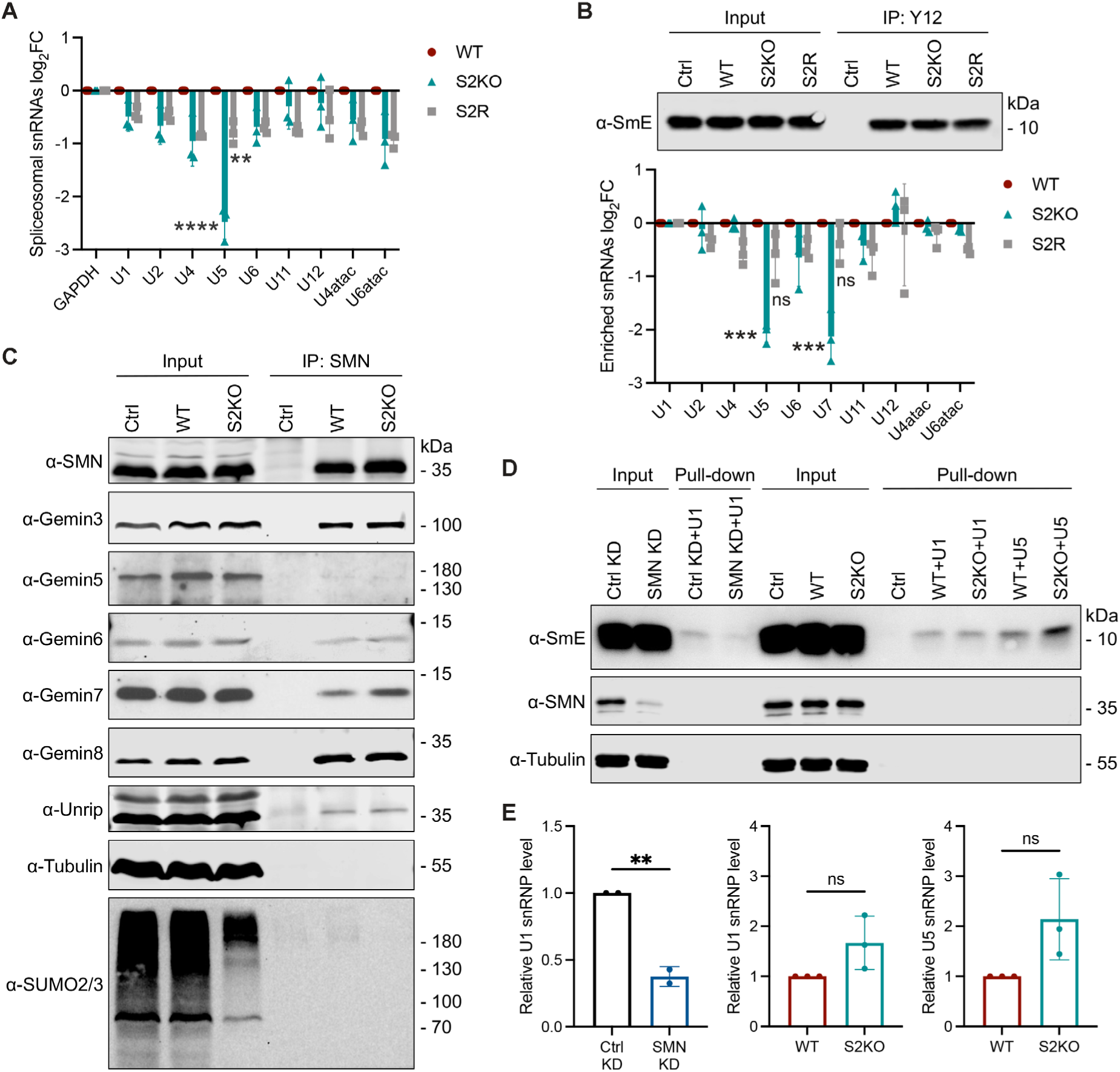
SUMO2 affects U7 snRNP levels independent of core SMN complex function. (A) RT-qPCR analysis of spliceosomal snRNAs using total RNA extracted from indicated cell lines from three independent replicates. *GAPDH* mRNA was used as an internal control. (B) Immunopurification of snRNP complexes with the Y12 antibody or IgG as control. Enriched snRNP complexes were analyzed by western blotting with antibodies against SmE. Enriched snRNAs from three independent replicates were analyzed by RT-qPCR and normalized to U1 snRNA. (C) Immunopurification of the SMN complex with antibodies against SMN. Subunits of the SMN complex were analyzed with western blotting. Control (Ctrl) condition was WT cell lysate incubated with IgG. (D) Western blot analysis of U1 and U5 snRNP in vitro assembly assays. U1 snRNP assembly in lysates from SMN-depleted (SMN KD) WT cells served as a positive control (first 4 lanes). Control knockdown (Ctrl KD) was WT cells treated with scrambled SMN siRNA oligos. Control (Ctrl) condition was WT cell lysate with no added snRNA. (E) Quantification of U1 and U5 snRNP assembly. Relative assembled snRNP levels were quantified from three independent replicates. *P* values were calculated using t tests (ns = not significant, * *p* < 0.05, ** *p* < 0.01, *** *p* < 0.001, **** *p* < 0.0001).

To investigate possible defects in SMN complex assembly, we immunopurified SMN complexes with antibodies against SMN, followed by western blotting with antibodies to individual protein subunits (Fig 6C, Fig S4A). This analysis revealed no defects in composition of the SMN complex in S2KO cells. To assess SMN complex activity, we performed in vitro snRNP assembly assays. In vitro-transcribed and biotinylated snRNAs were incubated with whole cell extracts and enriched with streptavidin-coated beads followed by western blotting with antibodies against SmE to quantify snRNP assembly activity (Fig 6D, E). We used SMN knockdown as a positive control and detected a ∼65% reduction in levels of assembled U1 snRNP compared to the control knockdown. We then measured U1 snRNP and U5 snRNP assembly in S2KO cell extracts and detected no significant differences in assembly of either snRNP compared to WT cell extracts. We were unable to detect U7 snRNP assembly due to exceedingly low levels.

We also analyzed SMN complex localization using immunofluorescence microscopy and detected no mislocalization to the cytoplasm as previously seen in cells depleted of the SUMO E2 conjugating enzyme Ubc9 (Riboldi et al. 2021) (Fig S4B). Our findings reveal reduction in U5 snRNP and U7 snRNP levels in S2KO cells but no detectable defects in the core SMN complex.

### A SUMO-interacting motif in Lsm11 facilitates U7 snRNP assembly

Previous studies identified Lsm11 as a SUMO2 substrate and mapped modification to lysine residues 120 and 260 (Fig 7A) (Hendriks et al. 2017). To assess whether SUMO2 affects U7 snRNP levels through modification of Lsm11 at these sites, we constructed a V5-tagged Lsm11 expression plasmid with lysine-to-arginine substitutions (K120/260R). We also constructed an expression plasmid for a V5-tagged SUMO2-Lsm11 fusion protein to mimic sumoylation (Ross et al. 2002; Kim et al. 2009). We transiently transfected these plasmids in U2OS cells and measured U7 snRNP expression levels as described above (Fig S5A). Transient over-expression of the Lsm11 K120/260R mutant in WT cells had no effect on U7 snRNP levels compared to wild-type Lsm11 over-expression. Similarly, over-expression of the SUMO2-Lsm11 fusion was unable to rescue U7 snRNP levels in S2KO cells.

**Figure 7.**
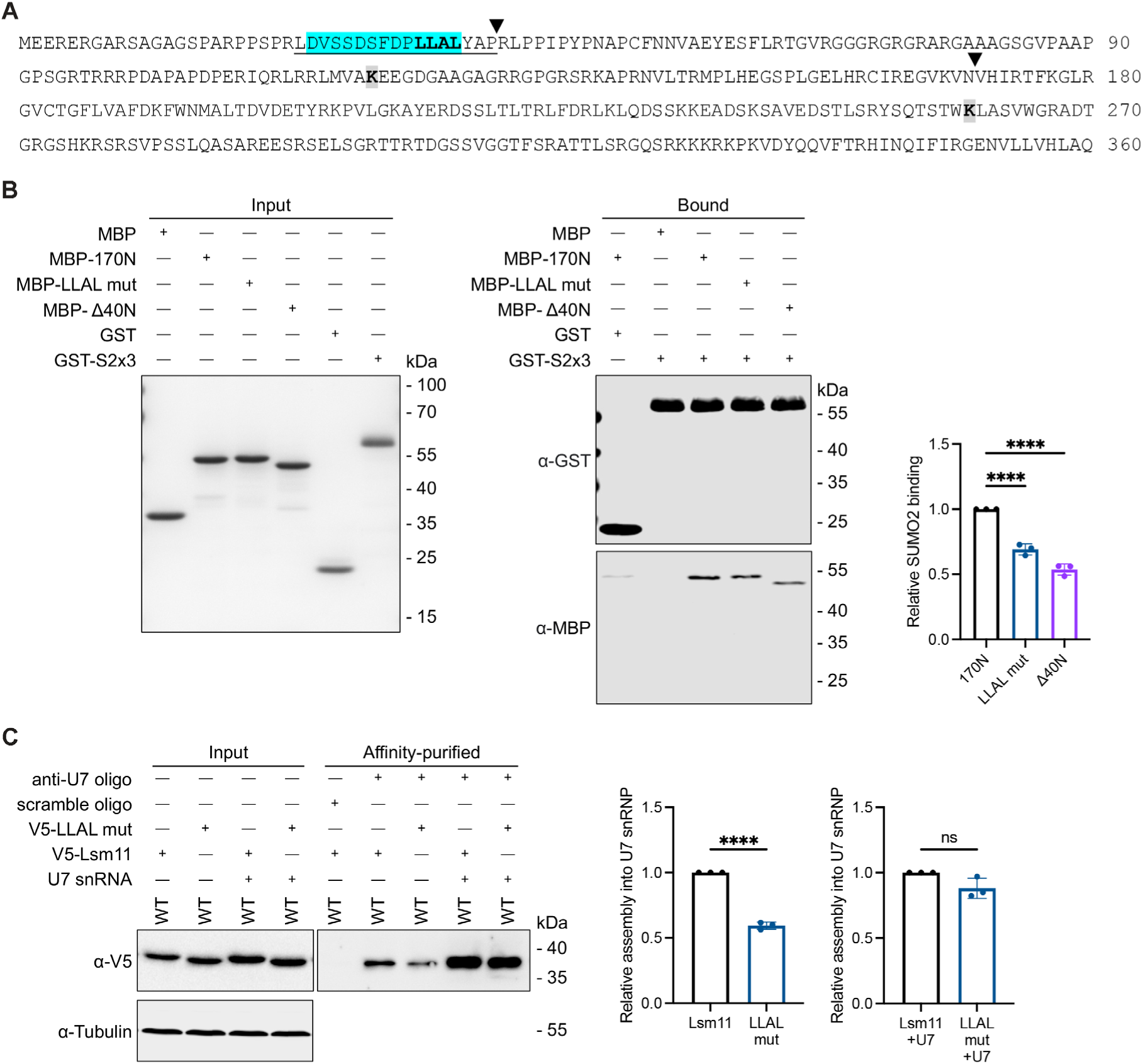
SUMO2 facilitates U7 snRNP assembly through direct binding to a SIM in the N-terminus of Lsm11. (A) Amino acid sequence of human Lsm11. Boundaries of the N-terminal 1-170 amino acid (170N) and 41-170 amino acid (Δ40N) fragments are indicated with arrowheads. The SIM is highlighted in blue, with the hydrophobic core (amino acids 34-37) in bold. Sumoylated lysines, K120 and K260, are in gray and bold. The FLASH-binding region is underlined. (B) In vitro SUMO2 binding assay. Purified GST-SUMO2 trimer (GST-S2x3) was bound to glutathione beads and incubated with the indicated MBP-Lsm11 N-terminal fragments. Purified GST and MBP proteins were used as controls. Inputs were analyzed with SimplyBlue staining and bound fractions were analyzed with western blotting. Relative SUMO2 binding was quantified from three independent replicates. (C) Western blot analysis of U7 snRNP complexes affinity-purified from WT cells transfected with V5-Lsm11 or V5-Lsm11 LLAL mutant (LLAL mut) individually, or co-transfected with U7 snRNA as indicated. Relative incorporation into U7 snRNP complexes was quantified from three independent replicates. *P* values were calculated using t tests or one-way ANOVA tests (ns = not significant, * *p* < 0.05, ** *p* < 0.01, *** *p* < 0.001, **** *p* < 0.0001).

In addition to containing covalent conjugation sites, we also identified a putative SIM within the N-terminus of Lsm11 between residues 25 and 37 (Fig 7A). To test whether Lsm11 binds to SUMO2, we performed in vitro SUMO2 binding assays (Fig 7B). We expressed and purified a GST-tagged polymeric SUMO2 trimer, and three MBP-tagged Lsm11 N-terminal fragments. The Lsm11-170N fragment contained amino acids 1-170. The Lsm11-Δ40N fragment comprised amino acids 41-170 and lacked the predicted SIM. The Lsm11-SIM mutant (LLAL mut) consisted of amino acids 1-170 and contained L34A, L35A, L37A mutations predicted to abolish SUMO2 binding (Hecker et al. 2006). GST-tagged trimeric SUMO2 was bound to glutathione-coated beads and subsequently incubated with MBP-tagged Lsm11 N-terminal fragments. Interactions were analyzed by western blotting (Fig 7B). Lsm11-170N showed direct binding to SUMO2, and this binding was reduced by ∼30% for the SIM mutant fragment and ∼50% for the SIM deletion fragment.

To further validate this interaction, we purified a GST-tagged trimeric SUMO2 QFI mutant, containing Q31A, F32A, and I34A mutations that disrupt SIM-binding, as well as an MBP-tagged Lsm10/Lsm11 heterodimer (Sun et al. 2007; Zhu et al. 2008). We again observed direct binding of the Lsm10/Lsm11 complex to SUMO2, and this binding was decreased for the SUMO2 QFI mutant (Fig S5B). Together, these results demonstrate that the N-terminus of Lsm11 contains a functional SIM.

To test whether the N-terminal SIM of Lsm11 functions in U7 snRNP assembly, we constructed a V5-tagged Lsm11 SIM mutant (LLAL mut) expression plasmid. We transiently over-expressed this mutant in U2OS WT cells and quantified its assembly into U7 snRNP complexes by affinity purification followed by western blotting (Fig 7C). The Lsm11 SIM mutant showed ∼50% reduction in incorporation into U7 snRNP complexes compared to wild-type Lsm11. This defect in assembly was rescued by co-transfection with U7 snRNA. These results reveal a functionally important interaction between the N-terminal SIM in Lsm11 and SUMO2 that enhances the efficiency of U7 snRNP assembly.

## DISCUSSION

In this study, we identified a novel function of SUMO2 in promoting canonical histone pre-mRNA processing through affecting FLASH dynamics and U7 snRNP assembly. Specifically, we found that SUMO2 stabilizes the association of FLASH with HLBs. We also found the interaction between a SIM in the N-terminus of Lsm11 and SUMO2 is critical for U7 snRNP expression levels, and that this interaction facilitates U7 snRNP assembly independent of core SMN complex functions. In the absence of Lsm11-SUMO2 interactions, U7 snRNP levels in the cell (and the HLB) are reduced, resulting in inefficient processing of histone pre-mRNAs.

### SUMO2 regulation of FLASH dynamics and HLB assembly

Previous studies have established a key role of SUMO in regulating PML-NB assembly. SUMO functions as “velcro” to drive the association of PML-NB proteins through non-covalent interactions between SUMO-modified proteins and proteins containing SIMs (Matunis et al. 2006; Shen et al. 2006). Consistent with this, we previously found that the numbers and size of PML-NBs were reduced in S2KO cells, indicating a paralog-specific function for SUMO2 in nuclear body formation and dynamics (Bouchard et al. 2021). In this study, we demonstrate that SUMO2 plays a role in HLBs, in part by stabilizing interactions between FLASH and other HLB-associated proteins. Sumoylation of FLASH at a lysine residue (K1813) has been shown to affect FLASH function and stability (Alm-Kristiansen et al. 2009; Vennemann and Hofmann 2013). FLASH also harbors four tandem SIMs in its C-terminus (Sun and Hunter 2012). miniFLASH contains both K1813 and one reported SIM (residues 1794-1798). Together with our findings, this suggests that SUMO2 modification and SUMO2-SIM interactions may affect the association of FLASH with HLBs. Other HLB-associated proteins, including Lsm11, symplekin, CPSF73, and NELF have also been identified as SUMO2 substrates (Vethantham et al. 2007; Hendriks and Vertegaal 2016; Hendriks et al. 2017; Gonzalez-Prieto et al. 2021; Rawat et al. 2021). This further suggests that, like PML-NBs, SUMO2-SIM interactions may play a role in the assembly and stabilization of HLBs. However, we did not observe a concentration of SUMO2 within HLBs, suggesting a more dynamic and transient function compared to PML-NBs (Fu et al. 2005; Keiten-Schmitz et al. 2021).

### SUMO2 is required for efficient U7 snRNP assembly

The U7 snRNP complex is present at only ∼1,000 copies per cell, and a reduction in its levels disrupts the coupling of canonical cleavage with transcription and leads to polyadenylation of histone mRNAs (Tisdale et al. 2013; Tatomer et al. 2016). Factors affecting U7 snRNP levels include SMN snRNP assembly activity and the levels of the U7 snRNP-specific Lsm10 and Lsm11 proteins. Notably, rescue of reduced U7 snRNP levels caused by SMN deficiencies requires over-expression of both Lsm10 and Lsm11 (Tisdale et al. 2022). Here, we found that rescue of U7 snRNP expression levels in S2KO cells was dependent only on over-expression of Lsm11 and U7 snRNA. These findings suggest that the reduced U7 snRNP levels in S2KO cells is likely due to U7 snRNP assembly defects distinct from those caused by SMN deficiencies. Consistent with this, we did not identify defects in composition of the SMN complex or spliceosomal snRNP assembly activity and spliceosomal snRNP levels were not affected, with the exception of U5 snRNP. Effects on U5 snRNP levels are likely due to specific roles of SUMO2 in regulating U5 snRNP biogenesis.

How does SUMO2 affect U7 snRNP assembly? We found that expressing a SUMO2-Lsm11 fusion protein or a sumoylation site mutant Lsm11 had no effects on U7 snRNP levels, suggesting that SUMO2 modification of Lsm11 is not required for U7 snRNP assembly. In contrast, we identified a consensus SIM in the N-terminus of Lsm11 that interacts with SUMO2. SUMO2 binding was not fully eliminated by deletion of this SIM, suggesting the presence of an additional non-consensus SIM in the N-terminus of Lsm11. In WT cells, expressing the SIM-mutant Lsm11 impaired the incorporation of Lsm11 into U7 snRNP complexes, and this defect was rescued by over-expressing U7 snRNA. Together, these results indicate that non-covalent interactions between the N-terminus of Lsm11 and SUMO2 promote U7 snRNP assembly. We interpret the rescue of U7 snRNP assembly by over-expression of Lsm11 and U7 snRNA to suggest two possible roles for SUMO2. First, SUMO2 may function to stabilize Lsm11 and maintain expression levels required for efficient U7 snRNP assembly. Alternatively, SUMO2 may function to enhance the interaction between Lsm11 and a factor required for U7 snRNP assembly.

Notably, the SIM we identified in Lsm11 is located within an unstructured N-terminal tail not found in other Sm proteins and essential for interactions with FLASH (Fig 7A) (Yang et al. 2011; Yang et al. 2013; Skrajna et al. 2016). The FLASH-binding region overlaps completely with the SIM, containing both the hydrophobic core and upstream negatively charged aspartic acid and serine residues (Song et al. 2004; Hecker et al. 2006; Cappadocia et al. 2015). This N-terminal tail may destabilize Lsm11 prior to its incorporation into the U7 snRNP and association with FLASH. One of the earliest steps in U7 snRNP assembly occurs during Lsm11 translation. Fully synthesized Lsm11 remains bound to ribosomes prior to heterodimerization with Lsm10 and subsequent association with the Sm ring assembly chaperone plCln (Paknia et al. 2016). These interactions are thought to ensure proper Lsm11 folding in the cytoplasm prior to ring assembly and U7 snRNA loading. Lsm11-SUMO2 interactions could occur during these early assembly stages in the cytoplasm and promote Lsm11 stability, either directly or by recruiting other protein chaperones. Consistent with this, roles for SUMO1 in protecting proteins from aggregation in the cytoplasm have been previously reported (Wang et al. 2023). Following assembly, the loading of Sm ring complexes onto spliceosomal snRNAs is dependent on snRNA recognition by the Gemin5 subunit of the SMN complex (Battle et al. 2006b; Wahl and Fischer 2016). Gemin5, however, does not recognize the U7 snRNA, and SMN specificity factors involved in loading U7 Sm rings onto U7 snRNA remain unknown. Thus, SUMO2 could also play a role at this step by facilitating interactions between Sm rings containing Lsm11 and specificity factors required for U7 snRNA recognition. The identification of SUMO2-modified factor(s) affecting U7 snRNP assembly will be essential to further exploring these possible functions.

### U7 snRNP localization to HLBs and interactions with processing factors

Following assembly in the cytoplasm, the U7 snRNP is imported into the nucleus and concentrated in HLBs. Though the mechanisms affecting its HLB localization are unclear, this process is independent of FLASH-Lsm11 interactions (Burch et al. 2011; Tatomer et al. 2016). Thus, interactions with SUMO2 may function to maintain Lsm11 and U7 snRNP stability prior to HLB localization. Within HLBs, Lsm11-SUMO2 interactions could also regulate the timely interaction with FLASH and activation of histone pre-mRNA processing specifically in S phase. Other functions are suggested by roles for SUMO in regulating mRNA polyadenylation through modification of CPSF73 and symplekin, factors shared with the histone cleavage complex (Vethantham et al. 2007).

## Conclusions

In summary, we favor two primary mechanisms by which SUMO2 may promote canonical histone pre-mRNA processing. First, SUMO2 can enhance the dynamic association and concentration of FLASH, and predictably other factors, within HLBs. Second, SUMO2 can facilitate the efficiency of U7 snRNP assembly through non-covalent interactions with the N-terminus of Lsm11, possibly through stabilizing Lsm11 proteins in the cytoplasm and the nucleus prior to association with FLASH, and enhancing interactions between Lsm11 and U7 snRNP-specific assembly factors. These proposed functions for SUMO2 are not mutually exclusive and are compatible with the known diversity of effects of sumoylation. Collectively, our studies define SUMO2 as an important component of the canonical histone pre-mRNA 3’ end processing pathway. Insights from these studies will contribute to a mechanistic understanding of the misregulation of canonical histone pre-mRNA processing that occurs under disease conditions.

## MATERIALS AND METHODS

### Ribo-minus RNA-Seq

Total RNA isolated using the Sigma-Aldrich GenElute Mammalian Total RNA Miniprep kit (RTN70) and sent to Novogene for ribo-minus selection and RNA sequencing. Using the TopHat2 software, the sequencing reads were processed by trimming and mapping them to the human hg38 reference genome (Kim et al. 2013). The maximum insertion and deletion length was set to 2 base pairs and default parameters were used for the remaining settings. In addition, only reads that were uniquely mapped, excluding any ’mixed’ or ’discordant’ reads from the results, were retained.

### Cell lines and cell culture conditions

U2OS WT, S1KO, S2KO, and S2R cell lines have been previously described and were grown at 37°C, 5% CO_2_ in DMEM (Gibco, 11965118) supplemented with 10% fetal bovine serum (Sigma-Aldrich, F2442) (Bouchard et al. 2021).

### Cell transfection

#### Plasmids and U7 snRNA

5 x 10^5^ cells were seeded and grown overnight. 1 μg of plasmid was transfected into the cells using Lipofectamine 2000 (Invitrogen, 11668027). 48 hr after transfection, cells were harvested for downstream analysis. When co-transfecting with U7 snRNA, 500 ng of capped U7 snRNA generated by in vitro transcription was transfected 24 hr after plasmid transfection using Lipofectamine RNAiMax (Invitrogen, 13778150). 24 hr after U7 snRNA transfection, cells were harvested for western blot analysis. When transfecting U7 snRNA in WT-V5-Lsm11 and S2KO-V5-Lsm11 stable cells, 2 x 10^4^ cells were seeded in a 6-well plate and grown overnight. 20 ng and 80 ng of in vitro-transcribed and capped U7 snRNA was transfected at 24 hr and 72 hr, respectively. Cells were collected 96 hr after seeding for downstream RT-qPCR analysis.

#### SMN siRNA knockdown

5 x 10^5^ cells were seeded and grown overnight. 20 nM of SMN siRNA oligo was transfected into the cells using Lipofectamine RNAiMax. 48 hr after transfection, cells were harvested for downstream analysis.

### FRAP

3 x 10^4^ cells were seeded in Nunc Lab-Tek II chambered coverglass slides (155382PK) and grown overnight. 175 ng of GFP-miniFLASH plasmid was transfected into the cells using Lipofectamine 2000. 24 hr after transfection, FRAP analysis was performed using a ZEISS LSM 900 confocal microscope. FRAP data were analyzed using FIJI software (Schindelin et al. 2012).

### RT-qPCR analysis of histone mRNA polyadenylation

These assays were performed as previously described (Bouchard et al. 2021). Experimental details used in this study are included in the supplemental material.

### HLB imaging and quantification

These assays were performed as previously described (Bouchard et al. 2021). Experimental details used in this study are included in the supplemental material.

### U7 snRNP affinity purification

These assays were performed as previously described (Smith et al. 1991). Experimental details used in this study are included in the supplemental material.

### Immunopurification of SMN and spliceosomal snRNPs

These assays were performed as previously described (Zhang et al. 2008). Experimental details used in this study are included in the supplemental material.

### snRNA RT-qPCR analysis

These assays were performed as previously described (Zhang et al. 2008). Experimental details used in this study are included in the supplemental material.

### In vitro transcription of snRNAs

These assays were performed as previously described (Boskovic et al. 2020). Experimental details used in this study are included in the supplemental material.

### In vitro snRNP assembly assays

These assays were performed as previously described (So et al. 2016). Experimental details used in this study are included in the supplemental material.

### SUMO2 and Lsm11 protein purification

These assays were performed as previously described (Zhu et al. 2008; Yang et al. 2023). Experimental details used in this study are included in the supplemental material.

## ACKNOWLEDGEMENTS

We thank Dr. Zbigniew Dominski (University of North Carolina at Chapel Hill) for the U7 antisense oligos and Lsm11 expression vectors and insightful advice. We thank Drs. Marcel Köhn and Stefan Hüttelmaier (Martin-Luther-University Halle-Wittenberg) for the miniFLASH plasmid and Dr. Livio Pellizzoni (Columbia University) for the Lsm10/11 co-expression plasmid and Gemin antibodies. We thank Dr. Francesco Lotti (Columbia University) for helpful suggestions on SMN. This work was supported by grants from the National Institutes of Health (GM060980, M.J.M., T32CA009110, C.M.M. and M.W.S., R35GM118093, L.T.) and a fellowship from the American Heart Association and the District of Columbia Women’s Board (https://doi.org/10.58275/AHA.24PRE1240229.pc.gr.190686, S.H.).

## Author contributions

S.H., M.J.M. conceptualization; S.H., M.W.S., A.D., C.M.M., W.W. experimentation; S.H., P.L., B.F. analysis and visualization; S.H., M.J.M. writing; J.Q., L.T., W.F.M., M.J.M. supervision.

